# Pleiotropic effect of ABCG2 in gout: involvement in serum urate levels and progression from hyperuricemia to gout

**DOI:** 10.1101/2020.01.06.894899

**Authors:** Rebekah Wrigley, Amanda J Phipps-Green, Ruth K Topless, Tanya J Major, Murray Cadzow, Philip Riches, Anne-Kathrin Tausche, Matthijs Janssen, Leo AB Joosten, Tim L Jansen, Alexander So, Jennie Harré Hindmarsh, Lisa K Stamp, Nicola Dalbeth, Tony R Merriman

## Abstract

**Background:** The ABCG2 Q141K (*rs2231142*) and *rs10011796* variants associate with hyperuricaemia (HU). The effect size of *ABCG2 rs2231142* on urate is ∼60% that of *SLC2A9*, yet the effect size on gout is greater. We tested the hypothesis that ABCG2 plays a role in the progression from HU to gout by testing for association of *ABCG2 rs2231142* and *rs10011796* with gout using HU controls.

**Methods:** We analysed 1,699 European gout cases and 14,350 normourciemic (NU) and HU controls, and 912 New Zealand (NZ) Polynesian (divided into Eastern and Western Polynesian) gout cases and 696 controls. Association testing was performed using logistic and linear regression with multivariate adjusting for confounding variables.

**Results:** In Europeans and Polynesians, the *ABCG2* 141K (T) allele was associated with gout using HU controls (OR=1.85, *P*=3.8E^-21^ and OR_meta_ =1.85, *P*=1.3E^-03^, respectively). There was evidence for an effect of 141K in determining HU in European (OR=1.56, *P*=1.7E^-18^) but not in Polynesian (OR_meta_=1.49, *P*=0.057). For *SLC2A9 rs11942223*, the T allele associated with gout in the presence of HU in European (OR=1.37, *P*=4.7E^-06^), however significantly weaker than *ABCG2 rs2231142* 141K (*P*_Het_=0.0023). In Western Polynesian and European, there was epistatic interaction between *ABCG2 rs2231142* and the genetically-independent *rs10011796*. Combining the presence of the 141K allele with the *rs10011796* CC-genotype increased gout risk, in the presence of HU, 21.5-fold in Western Polynesian (*P*=0.009) and 2.6-fold in European (*P*=9.9E^-06^). The 141K allele positively associated with flare frequency in Polynesian (*P*_meta_ *=*2.5E^-03^).

**Conclusion:** These data are consistent with a role for *ABCG2* 141K in gout in the presence of established HU.

## Introduction

The pathogenesis of gout is thought to require progression through three checkpoints: hyperuricaemia (HU), deposition of monosodium urate (MSU) crystals into articular and peri-articular structures, and an inflammatory response to these crystals (1). Genome-wide association studies (GWAS) have emphasised the contribution to urate control of genetic variation in renal and gut urate transporters, including *SLC2A9* and *ABCG2* (2, 3). When combined, variants in these two genes explain 3-4% of variance in urate levels and have strong effects on the risk of gout (4, 5). However, understanding of pathways regulating MSU crystal deposition and the inflammatory response to deposited crystals in gout remains important because fewer than a quarter of people with HU develop gout (6). Production of the inflammatory cytokine interleukin-1β (IL-1β) is central to the inflammatory response to MSU crystals (7, 8). The pathway that produces IL-1β involves activation of the NLRP3 inflammasome, resulting in cleavage of pro-IL-1β to mature IL-1β by caspase-1. There is little knowledge, however, about the genetic variants that promote the formation of MSU crystals and initiate the innate immune response in the presence of HU (9), although variants in the toll-like receptor 4 and components of the NLRP3 inflammasome have been associated with increased risk of gout (10-12).

The *ABCG2* gene was first associated with serum urate levels and gout by Dehghan et al (13). The encoded protein (also known as breast cancer resistance protein) functions as a urate and oxypurinol transporter in the kidney and gut (14). The lysine (T) allele of the Q141K (*rs2231142*) single nucleotide polymorphism (SNP) is associated with HU and increased risk of gout (reviewed in (14)). This variant decreases gut excretion of urate, contributing to a subtype of HU termed extra-renal under-excretion of urate (15, 16). The 141K allele is also associated with a poor response to the principal urate-lowering therapy allopurinol (xanthine oxidase inhibitor) in people with gout (17-19). Wen et al. reported a second genetically-independent variant, *rs10011796*, associated with poor allopurinol response, although this association could not be replicated in a well-phenotyped, allopurinol-compliant sample set (19). SNP *rs10011796* strongly associated with urate in Europeans (β=0.089 mg/dL, *P*=2.4×10^−51^), albeit with a weaker effect than *rs2231142* (β=0.221 mg/dL, *P*=4.4×10^−116^) (4).

The 141K variant creates instability in the protein’s nucleotide-binding domain, reducing expression of *ABCG2* due to a processing defect and impaired trafficking to the cell membrane (20). This causes a 50% reduction of ABCG2-mediated uric acid excretion (21). Dysfunction can be rescued by low temperature (22), and administration of small ligands, such as histone deacetylase inhibitors and colchicine, an anti-inflammatory drug used on the treatment of gout flares by disrupting neutrophil microtubule functioning (20, 23). The defective protein is retained in aggresomes, a cellular pathway activated when proteasome activity is exceeded, and is subsequently degraded by the autophagy pathway (23, 24). Deficiency in ABCG2 generates dysfunctional mitochondria (25) and reduced copy number of mitochondrial DNA associates with increased risk of gout in NZ Polynesian (26).

The observation that colchicine is able to rescue the 141K trafficking defect (23), the proposal that autophagy machinery and the inflammasome interact in the innate immune response (27), and evidence for association of *ABCG2 rs2231142* with gout in the presence of HU in East Asian populations (28-30), suggests that ABCG2 may be important in gout beyond its established role in elevating urate levels. This hypothesis is further supported by the observation that the effect size of *ABCG2* on urate in Europeans and Japanese is 58% and 73% that of *SLC2A9*, the most influential urate locus (4, 31), respectively, yet the effect size of *ABCG2* on gout is consistently larger than that of *SLC2A9* (4, 5, 32). Therefore we tested the hypothesis in European and Aotearoa New Zealand Polynesian (NZ Māori and Pacific Islands peoples) that *ABCG2* has a role in the progression of HU to gout using a genetic epidemiological approach by testing for association of *ABCG2 rs2231142* and *rs10011796* with gout in the presence of HU.

## Participants and Methods

### Participants

The European sample set comprised 1,699 participants with gout and 14,350 controls (2,422 asymptomatic HU, and 11,928 NU). The NZ Polynesian sample set (individuals of NZ and Cook Island Māori, Samoan, Tongan, Niuean and Tokelauan ancestry) comprised 912 participants with gout, and 696 controls (202 HU and 494 NU) (Additional File 1). Hyperuricemia was defined, for both sexes, as serum urate ≥0.42 mmol/L (7 mg/dL).

All people with gout fulfilled the 1977 American Rheumatism Association gout classification criteria (33). Gout cases were recruited from New Zealand (979 Europeans, 912 Polynesians), Australia and Europe (720 Europeans). The 14,350 European controls (all self-reported as not having physician-diagnosed gout, not having kidney disease and not taking urate-lowering medication) were obtained from five sources: 452 individuals recruited from New Zealand, 6,970 participants from the Atherosclerosis Risk in Communities (ARIC) study, 2,689 participants from the Framingham Heart Study (FHS), 1,492 participants from the Coronary Artery Risk Development in Young Adults (CARDIA) study, and 2,747 participants from the Cardiovascular Health Study (CHS). Phenotypes from baseline exams were used for all studies with the exception of CARDIA, where phenotypes from exam six were used. The 696 Polynesian controls were recruited from New Zealand using the same exclusion criteria.

Ancestry for controls from ARIC, FHS, CARDIA and CHS was used as provided in these datasets. Ancestry for the NZ, Australian and European gout cases and the NZ controls was classified based on principal components computed from genome-wide SNP data. The NZ Polynesian sample set was divided into Eastern and Western Polynesian ancestral groups based on principal components from genome-wide genotype data (34). Eastern Polynesian comprised of Cook Island and NZ Māori (543 cases and 462 controls), while Western Polynesian comprised of Samoa, Tonga, Tuvalu, Niue and Tokelau (369 cases and 234 controls) (35). A separate NZ Māori sample set of 124 cases and 50 controls is included within the Eastern Polynesian sample set data presented above. These participants were recruited in collaboration with Ngāti Porou Hauora Charitable Trust from the Ngāti Porou *rohe* (tribal territory) located in the East Coast (Tairāwhiti) region of the North Island of New Zealand.

### Genotyping

Three SNPs were examined: *rs2231142, rs10011796* (both *ABCG2*) and *rs11942223 (SLC2A9*). Genotypes for the NZ gout cases and controls, and for the European and Australian gout cases were determined using either a) Taqman^®^ assays (*rs2231142*: C_15854163_70; *rs10011796*: C_9510320_10; *rs11942223:* C_1216479_10; Applied Biosystems, Foster City, USA) on a Lightcycler^®^ 480 machine (Roche Applied Science, IN, USA), or b) Illumina Infinium CoreExome v24 bead chips processed at the University of Queensland (Centre for Clinical Genomics). Bead chip genotypes were auto-clustered using GenomeStudio v2011.1 software (Illumina, San Diego, USA). The Illumina GenomeStudio best practice guidelines and quality control protocols of Guo *et al.* (36) were applied to these auto-clustered genotypes to ensure final genotype calls were of the highest possible quality. SNP *rs7442295* was identified as being in complete linkage disequilibrium with *rs11942223* in European and East Asian populations using LDLink (37), and was used as a proxy on the bead chip for *rs11942223*. 2,940 samples were genotyped on both platforms for all 3 SNPs, with genotype concordance exceeding 99.8%.

Publicly-available genotype data for the ARIC, FHS, CARDIA and CHS participants were used. These genotypes were generated by the Affymetrix 6.0 platform (ARIC and CARDIA datasets), the combined Affymetrix 50K and 500K platform (the FHS dataset), and the Illumina Human CNV370v1 platform (the CHS dataset). *Rs2231142* was genotyped on the Affymetrix 50K and 500K platforms and imputed using IMPUTE2 using the 1000 Genomes Phase I haplotype reference panel in the ARIC, CARDIA and CHS sample sets. Similarly, *rs10011796 (ABCG2)* and *rs11942223* (*SLC2A9*) were imputed in all four sample sets.

### Statistical analysis

All analyses were performed using R v3.1.2 using RStudio 0.98. Deviation from Hardy-Weinberg equilibrium within each group (gout, HU controls and NU controls) was tested separately using the Haldane Exact test for Hardy-Weinberg equilibrium (α=0.01) for each population and each SNP. Allelic association testing of *ABCG2 rs2231142* and *rs10011796*, and *SLC2A9 rs11942223* for comparison, with gout and HU was performed using logistic regression. We analysed gout vs. all controls, NU controls vs. HU controls, and gout vs. HU controls. The SNPs were also tested for association with serum urate concentration at recruitment in combined NU and HU controls using linear regression. Association of each of the three variants with self-reported number of gout flares over a 12-month period was examined using linear regression. Analyses were adjusted by age, sex (except male-only analyses) and principal components (PCs) 1 to 10 (Polynesian sample sets only). For association testing of gout in the presence of HU, analyses were additionally adjusted by highest recorded serum urate concentration. All Polynesian meta-analyses were performed in R using meta v4.8-2 (38), and an inverse-variance fixed effect model unless otherwise stated.

Given the low linkage disequilibrium between *ABCG2 rs2231142* and *rs10011796* in European and East Asian populations (r^2^=0.087 and 0.12, respectively), genotypes were categorised as risk allele-positive or -negative for both SNPs, and association of *rs2231142-rs10011796* genotype combinations were tested for gout vs. all controls, and gout vs. HU controls using logistic regression. Non-additive interaction between the two SNPs in determining the risk of gout was also examined, including an interaction term between *rs2231142* and *rs10011796* in the multivariate-adjusted logistic regression model.

## Results

Relevant demographic, anthropomorphic and biochemical characteristics of the various sample sets are presented in Additional File 1. At recruitment urate levels were higher in the HU group than in the gout group in both European (0.47 vs 0.41mmol/L) and Polynesian (0.48 vs 0.43 mmol/L). In both European and Polynesian there was a preponderance of males in the gout (84.7 and 82.5 %, respectively) and HU sample sets (78.8 and 73.2 % respectively) compared to NU (40.0 and 36.9 %, respectively). Given this, along with evidence that *ABCG2 rs2231142* has a stronger effect on urate and gout in males than females (13), male-only analyses were also done (Additional File 2).

The genotypes of all sample sets were in Hardy-Weinberg equilibrium (*P*>0.01). *ABCG2* SNPs *rs2231142* (Q141K) and *rs10011796* and, for comparison, the *SLC2A9* SNP *rs11942223* were tested for allelic association with a) gout in the presence of HU (Table 1; gout vs. HU), b) gout *per se* (Table 1; gout vs. all controls), c) HU (Table 1; HU vs. NU) and, d) serum urate levels in controls (Table 2). In European, the T allele of *rs2231142* (141K) was strongly associated with HU using NU controls (OR=1.56, *P*=1.7E^-18^), and with gout when using HU controls (OR=1.85, *P*=3.8E^-21^). In the Polynesian sample sets, there was no evidence for an effect of *rs2231142* in determining HU in Western Polynesian (OR=1.52, *P*=0.091) or Eastern Polynesian (OR=1.41, *P*=0.37) (Table 1), with a Polynesian meta-analysis yielding OR_meta_=1.49, *P*=0.057. We note that the Polynesian ORs were similar to that in Europeans, suggesting that the non-significance may be due to reduced power of the Polynesian sample set. However, *rs2231142* was associated by linear regression with serum urate levels in Polynesian (Table 2; β_meta_=0.018 mmol/L, *P*=0.014) with an effect size similar to European (β=0.017 mmol/L, *P*=8.9E^-30^). In Polynesian, *rs2231142* was associated with the risk of gout compared to HU in Western Polynesian (OR=1.77, *P*=0.012), EP (OR=2.02, *P*=0.043), and the Polynesian meta-analysis (OR_meta_ =1.85, *P*=1.3E^-03^) with effect sizes similar to that in Europeans (Table 1).

**Table 1.** Association analysis of *rs2231142, rs10011796 (ABCG2)* and *rs11942223 (SLC2A9)* in European and NZ Polynesian samples sets

| SNP | Raw N | Genotype, N (%)<sup>†</sup> |  |  | Effect allele, N (%) | Gout vs. HU*<sup>^</sup><br>OR (95% CI), *P* | Gout vs. All Controls*<br>OR (95% CI), *P* | HU vs. NU*<br>OR, 95% CI, *P* |
| --- | --- | --- | --- | --- | --- | --- | --- | --- |
| ***rs2231142*** |  | **GG** | **GT** | **TT** | **T** |  |  |  |
| European gout | 1637 | 1006 (61.5) | 551 (33.7) | 80 (7.5) | 711 (21.7) | 1.81 (1.60-2.06), *3.0E-20* |  | - |
| European HU | 2378 | 1762 (74.1) | 577 (24.3) | 39 (1.6) | 655 (13.8) | 1.85 (1.63-2.11), *3.8E-21* | 2.42 (2.18-2.68), *1.9E-64* | 1.56 (1.41-1.72), *1.7E-18* |
| European NU | 11604 | 9428 (81.2) | 2053 (17.7) | 123 (1.1) | 2299 (9.1) |  |  |  |
| WP<sup>#</sup> gout | 369 | 96 (26.0) | 189 (51.2) | 84 (22.8) | 357 (48.4) | 1.96 (1.30-3.00), *1.6E-03* |  | - |
| WP HU | 88 | 48 (54.6) | 29 (33.0) | 11 (12.5) | 51 (29.0) | 1.77 (1.14-2.80), *0.012*<sup>‡</sup> | 2.36 (1.73-3.26), *1.1E-07* | 1.52 (0.94-2.50), *0.091*<sup>‡</sup> |
| WP NU | 146 | 84 (57.5) | 52 (35.6) | 10 (6.9) | 72 (24.7) | - |  |  |
| EP<sup>#</sup> gout | 543 | 436 (80.3) | 106 (19.5) | 1 (0.2) | 108 (9.9) | 1.99 (1.05-4.05), *0.044* |  | - |
| EP HU | 114 | 101 (88.6) | 13 (11.4) | 0 (0.0) | 13 (5.7) | 2.02 (1.05-4.16) *0.043*<sup>‡</sup> | 2.54 (1.66-3.97), *3.4E-05* | 1.41 (0.65-2.93), *0.37*<sup>‡</sup> |
| EP NU | 348 | 315 (90.5) | 32 (9.2) | 1 (0.3) | 34 (4.9) |  |  |  |
| ***rs10011796*** |  | **CC** | **CT** | **TT** | **T** |  |  |  |
| European gout | 1580 | 339 (21.5) | 764 (48.4) | 477 (30.2) | 1718 (54.4) | 1.39 (1.26-1.53), *2.0E-11* | 1.55 (1.44-1.69), *1.3E-27* | - |
| European HU | 2422 | 666 (27.5) | 1208 (49.9) | 548 (22.6) | 2304 (47.6) | 1.39 (1.26-1.53), *3.5E-11* |  | 1.18 (1.10-1.26), *7.8E-07* |
| European NU | 11927 | 3688 (30.9) | 5986 (50.2) | 2253 (18.9) | 10492 (44.0) |  | - |  |
| WP gout | 369 | 38 (10.3) | 163 (44.2) | 168 (45.5) | 499 (67.6) | 1.28 (0.86-1.91), *0.22* |  | - |
| WP HU | 88 | 10 (11.4) | 43 (48.9) | 35 (39.8) | 113 (64.2) | 1.27 (0.83-1.94), *0.27* | 1.32 (0.99-1.77), *0.062* | 1.02 (0.66-1.60), *0.92* |
| WP NU | 146 | 21 (14.4) | 67 (45.9) | 58 (39.3) | 183 (62.7) |  |  |  |
| EP gout | 543 | 93 (17.2) | 259 (47.9) | 189 (34.9) | 637 (58.9) | 0.94 (0.68-1.29), *0.69* |  | - |
| EP HU | 114 | 18 (15.8) | 51 (44.7) | 45 (39.5) | 141 (61.8) | 0.92 (0.66-1.27), *0.60* | 1.03 (0.83-1.27), *0.80* | 1.13 (0.81-1.57), *0.48* |
| EP NU | 348 | 67 (19.3) | 173 (49.7) | 108 (31.0) | 389 (55.9) |  |  |  |
| ***rs11942223*** |  | **TT** | **TC** | **CC** | **C** |  |  |  |
| European gout | 1576 | 1168 (74.1) | 382 (24.2) | 26 (1.6) | 434 (13.8) | 0.72 (0.63-0.83), *2.9E-06* |  | - |
| European HU | 2422 | 1658 (68.5) | 701 (28.9) | 63 (2.6) | 827 (17.1) | 0.73 (0.63-0.83), *4.7E-06* | 0.55 (0.49-0.62), *1.4E-25* | 0.66 (0.61-0.73), *2.2E-21* |
| European NU | 11927 | 7101 (59.5) | 4220 (35.4) | 606 (5.1) | 5432 (22.8) |  |  |  |
| WP gout | 369 | 351 (95.1) | 18 (4.9) | 0 (0.0) | 18 (2.4) | 0.53 (0.17-2.05), *0.31* | 0.61 (0.26-1.46), *0.26* | - |
| WP HU | 88 | 84 (95.5) | 4 (4.6) | 0 (0.0) | 4 (2.3) | 0.51 (0.15-2.08), 0.31 |  |  |
| WP NU | 146 | 135 (92.5) | 11 (7.5) | 0 (0.0) | 11 (3.8) |  |  | 0.83 (0.20-2.94), 0.78 |
| EP gout | 543 | 501 (92.3) | 39 (7.2) | 3 (0.6) | 45 (4.1) | 1.04 (0.47-2.61), 0.92 |  | - |
| EP HU | 114 | 105 (92.1) | 9 (7.9) | 0 (0.0) | 9 (3.9) | 1.24 (0.54-3.23), 0.64 | 1.06 (0.62-1.84), 0.84 |  |
| EP NU | 348 | 311 (89.4) | 37 (10.6) | 0 (0.0) | 37 (5.3) |  |  | 1.26 (0.51-12.94), 0.60 |

\*European sample set adjusted by age and sex. Polynesian sample sets additionally adjusted by PCs 1-10.
^For Gout vs. HU analyses, the top data is before adjustment by highest recorded serum urate, and the bottom figure is after adjustment.
†All sample sets are in Hardy-Weinberg equilibrium ( $P > 0.01$ ).
‡Polynesian meta-analysis of WP and EP Gout vs. HU, and of HU vs. NU yielded $OR_{meta}=1.85$ , $P=1.3E^{-03}$ , and $OR_{meta}=1.49$ , $P=0.057$ , respectively.
#WP, Western Polynesian; EP, Eastern Polynesian.

**Table 2.** Association analysis of *rs2231142, rs10011796 (ABCG2)* and *rs11942223 (SLC2A9)* with serum urate at recruitment (mmol/L) in European and NZ Polynesian controls

| SNP / Allele | N | Combined Sex*<br>$\beta$ mmol/L (SE) | P | N | Male Only*<br>$\beta$ mmol/L (SE) | P |
| --- | --- | --- | --- | --- | --- | --- |
| ***rs2231142* / T** |  |  |  |  |  |  |
| European control | 13982 | 0.017 (1.5E-03) | *8.9E-30* | 6514 | 0.020 (2.2E-03) | *5.7E-20* |
| WP control | 234 | 0.019 (9.2E-03) | *0.035* | 134 | 0.027 (0.011) | *0.017* |
| EP control | 462 | 0.015 (0.012) | *0.19* | 190 | 0.009 (0.023) | *0.68* |
| Polynesian meta-analysis | 696 | *0.018 (7.2E-03)* | *0.014* | 324 | *0.024 (0.010)* | *0.019* |
| ***rs10011796* / T** |  |  |  |  |  |  |
| European control | 14349 | 0.006 (9.2E-04) | *1.7E-10* | 6682 | 0.006 (4.7E-03) | *1.5E-06* |
| WP control | 234 | 0.006 (8.5E-03) | *0.51* | 134 | 0.006 (0.011) | *0.62* |
| EP control | 462 | 0.005 (5.0E-03) | *0.33* | 190 | 0.000 (8.1E-03) | *0.99* |
| Polynesian meta-analysis | 696 | *0.005 (4.3E-03)* | *0.94* | 324 | *0.002 (6.6E-03)* | *0.77* |
| ***rs11942223* / C** |  |  |  |  |  |  |
| European control | 14349 | - 0.024 (1.1E-03) | *8.4E-106* | 6681 | - 0.017 (1.6E-03) | *8.1E-27* |
| WP control | 234 | - 0.021 (0.023) | *0.38* | 134 | - 0.038 (0.038) | *0.32* |
| EP control | 462 | - 0.008 (0.012) | *0.53* | 190 | 0.052 (0.022) | *0.022* |
| Polynesian meta-analysis<sup>^</sup> | 696 | - 0.011 (0.011) | *0.63* | 324 | *0.012 (0.044)* | *0.78* |

\*European sample set adjusted by age and sex (combined sex only). Polynesian sample sets additionally adjusted by PCs 1-10.
<sup>^</sup>As the test for heterogeneity between Western Polynesian (WP) and Eastern Polynesian (EP) was significant (p=0.043), a random effects model was used.

The strength of association of *ABCG2 rs2231142* with gout in the presence of HU in European was significantly different to *SLC2A9 rs11942223* (OR_risk allele_=1.85 for *rs2231142* compared to OR_risk allele_=1.37 for *rs11942223, P*_Het_ =2.1E^-03^). In contrast, the effect sizes on HU in Polynesian were not significantly different (OR_risk allele_=1.56 for *rs2231142* compared to OR_risk allele_=1.52 for *rs11942223, P*_Het_=0.64). In the Polynesian sample set, the minor allele of *rs11942223* was uncommon (<5% in most groups) with no nominally significant (*P*<0.05) associations detected in the three gout and HU analyses (Table 1). Male-only analysis yielded similar data (Additional File 2).

*ABCG2 rs10011796* displayed a similar pattern of association in Europeans as *rs2231142*, albeit with a weaker effect size (OR=1.39, *P*=3.5E^-11^ for gout vs. HU, and OR=1.18, *P*=7.8E^-07^ for HU vs. NU). However, *rs10011796* was not significantly associated in any of the three comparisons (gout vs. HU, gout vs. all controls, HU vs. NU) in any of the Polynesian sample sets (all P>0.06) (Table 1).

The three SNPs of interest were tested for association with gout flare frequency in each sample set (Table 3). The *ABCG2 rs2231142* T allele was associated with an additional 2.54 and 2.16 self-reported flares per year in the WP and meta-analysed Polynesian sample sets (Table 3; *P*=8.9E^-03^ and *P*_mate_ =2.5E^-03^, respectively) but not in European (*P*=0.97). Neither *rs10011796* nor *rs11942223* associated with flare frequency in any sample set.

**Table 3.** Association analysis of *rs2231142, rs10011796 (ABCG2)* and *rs11942223 (SLC2A9)* gout risk alleles with the number of self-reported gout flares in the previous year by gout patients

| SNP / Risk Allele | N | $\beta$ (SE)* | P |
| --- | --- | --- | --- |
| ***rs2231142* / T** |  |  |  |
| European gout | 921 | - 0.019 (0.479) | 0.97 |
| Western Polynesian gout | 359 | 2.54 (0.965) | 8.9E-03 |
| Eastern Polynesian gout | 481 | 1.69 (1.061) | 0.11 |
| *Polynesian meta-analysis* | 840 | 2.16 (0.714) | 2.5E-03 |
| ***rs10011796* / T** |  |  |  |
| European gout | 920 | 0.29 (0.396) | 0.47 |
| Western Polynesian gout | 359 | 0.46 (0.982) | 0.64 |
| Eastern Polynesian gout | 481 | -1.02 (0.638) | 0.11 |
| *Polynesian meta-analysis* | 840 | - 0.58 (0.535) | 0.28 |
| ***rs11942223* / T** |  |  |  |
| European gout | 924 | - 0.11 (0.596) | 0.85 |
| Western Polynesian gout | 359 | 1.80 (3.122) | 0.56 |
| Eastern Polynesian gout | 481 | 2.53 (1.440) | 0.080 |
| *Polynesian meta-analysis* | 840 | 2.40 (1.308) | 0.067 |

\*European sample set adjusted by age and sex. Polynesian sample sets additionally adjusted by PCs 1-10.

Non-additive interaction between *rs2231142* and *rs10011796* in determining the risk of gout was examined for gout vs. all controls, and gout vs. HU (Table 4). The interaction of *rs2231142* and *rs10011796* was significant in European gout vs. all controls (*P*=7.9E^-03^) but did not reach significance in the gout vs. HU analysis (*P*=0.062). There was no evidence for significant interaction in any of the Polynesian sample sets. Stratification by genotype groups (Table 5) revealed that the interaction was driven by a non-additive contribution to the risk of gout (gout vs. all controls) when the *rs2231142* risk-positive genotype group (GT,TT) was combined with the *rs10011796* risk-negative genotype (CC). The ORs for this genotype combination were higher than the ORs for risk-positive genotypes at both SNPs (European OR=3.78 vs. 3.37; Western Polynesian OR=4.81 vs. 4.40; Eastern Polynesian OR=2.88 vs. 2.45). The gout vs. HU controls analysis yielded similar findings in Europeans as using all controls, but with reduced ORs (Table 5). However, there was a strikingly high risk for gout in the *rs2231142* risk positive and *rs10011796* risk negative category in Western Polynesian (OR=21.5, *P*=8.6×10^−3^).

**Table 4.** Interaction terms between *rs2231142* and *rs10011796* in determining risk of gout in European and NZ Polynesian sample sets

| Sample Set | Gout vs. All Controls* |  |  | Gout vs. HU* |  |  |
| --- | --- | --- | --- | --- | --- | --- |
|  | N | OR<sub>interaction</sub> (95% CI) | P | N | OR<sub>interaction</sub> (95% CI) | P |
| European | 15559 | 0.61 (0.43-0.88) | *7.9E-03* | 3956 | 0.65 (0.41-1.02) | *0.062* |
| Western Polynesian | 603 | 0.62 (0.17-2.11) | *0.45* | 457 | 0.10 (0.01-1.09) | *0.059* |
| Eastern Polynesian | 1003 | 0.89 (0.25-2.92) | *0.85* | 655 | 1.48 (0.18-8.32) | *0.67* |
| *Polynesian meta-analysis* | *1606* | *0.75 (0.31-1.78)* | *0.51* | *1112* | *0.54 (0.13-2.32)* | *0.41* |

\*European sample set adjusted by age and sex. Polynesian sample sets additionally adjusted by PCs 1-10.

**Table 5.** Risk of gout in *rs2231142-rs10011796* genotype combinations classified by absence/presence of gout risk alleles

| Sample Set | Genotype |  | Gout vs. All Controls* |  |  |  | Gout vs. HU Controls* |  |  |  |
| --- | --- | --- | --- | --- | --- | --- | --- | --- | --- | --- |
|  | *rs2231142*<sup>^</sup> | *rs10011796*<sup>^</sup> | Controls<br>N (%) | Gout<br>N (%) | OR (95% CI) | *P* | HU Controls<br>N (%) | Gout<br>N (%) | OR (95% CI) | *P* |
| European |  |  | 13981 | 1578 |  |  | 2378 | 1578 |  |  |
|  | GG | CC | 3996 (28.6) | 280 (17.7) | 1.00 | - | 601 (25.3) | 280 (17.7) | 1.00 | - |
|  | GG | **CT, TT** | 7193 (51.4) | 687 (43.5) | 1.46 (1.26-1.71) | *9.2E-07* | 1161 (48.8) | 687 (43.5) | 1.33 (1.11-1.59) | *2.1E-03* |
|  | **GT, TT** | CC | 247 (1.8) | 59 (3.7) | 3.78 (2.67-5.28) | *1.8E-14* | 53 (2.2) | 59 (3.7) | 2.60 (1.70-3.98) | *9.9E-06* |
|  | **GT, TT** | **CT, TT** | 2545 (18.2) | 552 (35.0) | 3.37 (2.87-3.98) | *2.8E-48* | 563 (23.7) | 552 (35.0) | 2.24 (1.85-2.73) | *5.8E-16* |
| WP |  |  | 234 | 369 |  |  | 88 | 369 |  |  |
|  | GG | CC | 22 (9.4) | 14 (3.8) | 1.00 | - | 9 (10.2) | 14 (3.8) | 1.00 |  |
|  | GG | **CT, TT** | 110 (47.0) | 82 (22.2) | 1.47 (0.63-3.54) | *0.38* | 39 (44.3) | 82 (22.2) | 2.39 (0.81-6.98) | *0.11* |
|  | **GT, TT** | CC | 9 (3.8) | 24 (6.5) | 4.81 (1.53-16.06) | *8.5E-03* | 1 (1.1) | 24 (6.5) | 21.52 (3.05-449.83) | *8.6E-03* |
|  | **GT, TT** | **CT, TT** | 93 (39.7) | 249 (67.5) | 4.40 (1.92-10.33) | *5.1E-04* | 39 (44.3) | 249 (67.5) | 5.26 (1.84-14.74) | *1.6E-03* |
| EP |  |  | 462 | 541 |  |  | 114 | 541 |  |  |
|  | GG | CC | 77 (16.7) | 72 (13.3) | 1.00 | - | 16 (14.0) | 72 (13.3) | 1.00 | - |
|  | GG | **CT, TT** | 339 (73.4) | 362 (66.9) | 0.96 (0.63-1.46) | *0.85* | 85 (74.6) | 362 (66.9) | 0.83 (0.42-1.57) | *0.58* |
|  | **GT, TT** | CC | 8 (1.7) | 21 (3.9) | 2.88 (0.98-9.25) | *0.064* | 2 (1.8) | 21 (3.9) | 1.44 (0.31-10.58) | *0.67* |
|  | **GT, TT** | **CT, TT** | 38 (8.2) | 86 (15.9) | 2.45 (1.36-4.47) | *3.2E-03* | 11 (9.6) | 86 (15.9) | 1.78 (0.72-4.52) | *0.22* |

\*European sample set adjusted by age and sex. Polynesian sample sets additionally adjusted by PCs 1-10.
<sup>^</sup>The T allele (presence highlighted in bold) is the gout risk allele in European for both SNPs.

## Discussion

We report association of the *ABCG2 rs2231142* 141K (T) allele with gout in the presence of HU in European, Eastern and Western Polynesian sample sets. In all sample sets, the effect size was substantial (OR=1.77-2.02) (Table 1). These results are consistent with a role for *ABCG2* in the progression from HU to gout. This conclusion is supported by the association of 141K with flare frequency in Polynesians (*P*_meta_ =2.5E^-03^) (Table 3). For *rs10011796*, there was evidence for association with gout in the presence of HU and with HU in Europeans only. Stratification of genotypes by presence or absence of gout-risk allele in the HU vs gout analysis provides evidence for an epistatic interaction between *rs2231142* and *rs10011796* most notably in Western Polynesian (Table 5). The addition of the 141K risk allele to the non-risk *rs10011796* CC-genotype increased risk of gout for these individuals non-additively from 1.0 to 21.5 in the gout vs. HU analysis. We did observe an effect also for *SLC2A9 rs11942223* in the gout vs HU comparison for Europeans (OR=1.4 for risk allele), however this was notably weaker than that for *ABCG2 rs2231142* (OR=1.9; *P*_Het_=0.0023).

It is increasingly clear that there is heterogeneity in the frequency of the pathogenic 141K variant between the ancestrally-defined Western Polynesian (Samoa, Tonga, Niue, Tokelau) and Eastern Polynesian (New Zealand Māori, Cook Island Māori) populations of New Zealand. Although the T allele has a similar gout risk effect size, it is 5-fold more prevalent in Western Polynesian (Table 1) and will, therefore, have a greater impact on this population. It is also a known risk factor for tophus in the presence of gout in Western Polynesian (OR=1.66) but not in Eastern Polynesian (OR=0.91) (39). The interaction of the risk T allele of *rs2231142* with *rs10011796* in promoting gout in the presence of HU is observed in Western Polynesian but not Eastern Polynesian sample sets (Table 5). Finally, we found association of the 141K allele with flare frequency in Western Polynesian but not Eastern Polynesian (Table 3). Collectively, at *ABCG2* at least, these findings emphasize the need to carefully account for ancestry in studies investigating the genetic causes of gout in the Polynesian populations of New Zealand.

There are parallels between ABCG2 141K and the important cystic fibrosis-causing gene variant, ΔF508 in CFTR, also an ABC transporter. Both variants cause instability in the nucleotide binding domain of their respective proteins and can be corrected by small molecules (20, 23). Accumulation of aggresomes is a feature of both ABCG2 141K and cystic fibrosis (40), indicative of impaired and/or inadequate protein degradation. Dysfunctional autophagosome clearance in cystic fibrosis leads to a hyper-inflammatory state in which there is increased reactive oxygen species and impaired autophagy. This activates the NLRP3 inflammasome (41). In addition, there is accumulation of p62, a protein that regulates aggresome formation by delivering ubiquitinated proteins for degradation by autophagy, resulting in increased IL-1β production by promoting cleavage of pro-caspase-1 to caspase-1 (42). The IL-1β produced further increases p62 levels (40) resulting in defective autophagy in cystic fibrosis via accumulation of misfolded proteins in aggresomes. Reducing p62 levels allows localization of ΔF508 CFTR to the cell surface where it can function (40). In gout, MSU crystals impair proteasomal degradation causing increased expression of p62 (42), and it is likely that proteasomal degradation is impaired in the presence of the 141K variant as evidenced by the formation of aggresomes (23). ABCG2 promotes autophagy in cancer cell lines exposed to stressors such as nutrient deprivation (43), although the ability of the 141K allele to impair autophagy is not yet established. It is possible that defective autophagy resulting from the 141K variant could lead to increased IL-1β signaling, since autophagy normally allows for negative feedback regulation of IL-1β production via degradation of the NLRP3 inflammasome (44). Autophagy is necessary for formation of neutrophil extracellular traps that attenuate the inflammatory response to MSU crystals (45). Future studies that investigate and compare IL-1β production in response to MSU and autophagy in cells with the different 141Q and 141K alleles would be illuminating.

The molecular mechanism driving the non-additive interaction between *ABCG2* SNPs *rs2231142* and *rs10011796* is unclear. Given that *rs10011796* is a noncoding intronic variant outside any known Encode regulatory motif features (www.encodeproject.org), it is likely that *rs10011796* is in linkage disequilibrium with the causal variant, rather than the causal variant itself, although other intronic variants in *ABCG2* have been associated with ABCG2 expression (46, 47). The majority of common phenotype associations identified by GWAS are expression quantitative trait loci (48), influencing gene expression and transcript stability, and these variants can therefore modify the penetrance of coding variants (49). Thus, it is reasonable to hypothesise that *rs10011796* marks an effect that influences gene expression. This is consistent with the presence of an expression QTL (regulatory) effect independent of *rs2231142* in the *ABCG2* urate GWAS signal (47). The *rs10011796* C-allele (that associates with reduced serum urate and risk of gout) does associate with reduced *ABCG2* and increased *PPM1K-DT* (a long non-coding RNA 100kb downstream of *ABCG2*) expression (www.gtex.org). How *rs10011796* (or more likely a variant in linkage disequilibrium) could synergise with *rs2231142* to amplify the risk of gout is unclear. However it has previously been reported that a urate-associated variant at the *MAF* locus influences the expression of MAF via a long non-coding RNA (50). It is possible that in Western Polynesian people with HU, the combination of the 141K risk allele with the *rs10011796* CC-genotype has an epistatic effect where an altered amount of 141K is internalized, disrupting important stoichiometric relationships and promoting gouty inflammation. Finally, it is interesting to note that local epistatic interactions have also been reported at *SLC2A9* in the control of urate levels (51).

In addition to the common Q141K variant there are numerous other uncommon and rare missense variants in ABCG2, mostly detected by resequencing *ABCG2* exons in people with gout. These variants tend, as does 141K, to reduce the urate transport ability of ABCG2 (52), they associate with gout (52,53) and, including 141K, associate with an earlier age-of-onset of gout (52, 54). The effect size on gout is similar to that of 141K – increasing risk 2-3 fold (53). It is possible that the rare and uncommon variants also contribute to the progression from HU to gout, however testing this hypothesis will require very large datasets. Of significance for the study of the pathogenesis of gout, is the possibility that genetic variation in other genes contribute both to HU and the progression from HU to gout. In our data there was a suggestion that *SLC2A9* could be one such gene.

## Conclusion

We provide genetic epidemiological evidence supporting a role for ABCG2 141K in the progression from HU to gout, additional to its role in promoting HU. The variant may promote a hyper-inflammatory state akin to that observed with the cystic fibrosis gene, *CFTR* ΔF508, featuring defective autophagy, formation of aggresomes, and activation of the NLPR3 inflammasome.

## Abbreviations

ABCG2: ATP-binding cassette subfamily G member 2;
ARIC: Atherosclerosis Risk in Communities;
CARDIA: Coronary Artery Risk Development in Young Adults;
CTFR: cystic fibrosis transmembrane conductance regulator;
CHS: Cardiovascular Health Study;
EP: Eastern Polynesian;
FHS: Framingham Heart Study;
GWAS: genome-wide association study;
HU: hyperuricemia;
IL-1β: interleukin-1β;
MSU: monosodium urate;
NLRP3: NLR family pyrin domain containing 3;
NU: normouricemia;
NZ: New Zealand;
OR: odds ratio;
PC: principal component;
SLC2A9: solute carrier family 2 member 9;
SNP: single nucleotide polymorphism;
WP: Western Polynesian.

## Declarations

### Ethics approval and consent to participate

In New Zealand the New Zealand Multi-Region Ethics Committee (MEC/05/10/130) and the Northern Y Region Health Research Ethics Committee (Ngāti Porou Hauora Charitable Trust study; NTY07/07/074) provided ethical approval for the study. The following institutional committees in Europe and Australia also granted ethical approval: Research Ethics Committee, University of New South Wales; Ethikkommission, Technische Universität Dresden (EK 8012012); South East Scotland Research Ethics Committee (04/S1102/41); Commission Cantonale D’éthique de la Recherche sur l’être Humain, Université de Lausanne; Commissie Mensgebonden Onderzoek regio Arnhem - Nijmegen. All subjects gave written informed consent. The Database of Genotype and Phenotype (www.ncbi.nlm.nih.gov/gap) approval number was #834 for accessing data from the ARIC, FHS, CARDIA and CHS studies.

### Consent for publication

Not applicable.

### Availability of data and materials

Owing to consent restrictions it is not possible to make the New Zealand, Australian and European gout-control datasets publicly available, although they may be able to be made available from the corresponding author under appropriate request. The ARIC, FHS, CHS, CARDIA datasets are publicly-available at the Database of Genotype and Phenotype.

### Competing interests

The authors declare that they have no competing interests.

## Funding

The study was funded by the Health Research Council of New Zealand (grant 14/527). The funder had no role in the design, execution and reporting of the study.

## Author contributions

RW, AJP, TRM contributed to study design, analysed and interpreted the data and drafted the manuscript. RKT, TJM, MC contributed to data analysis, interpretation and the manuscript. PR, A-KT, MJ, LABJ, TLJ, AS, JHH, LKT, ND contributed to data collection, interpretation and contributed to the manuscript draft. All authors read and approved the final manuscript.

## Acknowledgements

The authors would like to thank Jordyn Allan, Jill Drake, Roddi Laurence, Christopher Franklin, Meaghan House and Gabrielle Sexton for recruitment. Matthew Brown, Linda Bradbury and The Arthritis Genomics Recruitment Initiative in Australia network are acknowledged. The European Crystal Network (55), formed after the first European Crystal Workshop in Paris, March 2010 (Prof Frédéric Lioté, Paris, and Prof Alexander So, Lausanne, convenors) is also acknowledged. The Atherosclerosis Risk in Communities and Framingham Heart study analyses (project #834) were approved by the relevant Database of Genotype and Phenotype (dbGaP; www.ncbi.nim.nih/gov/dbgap) Data Access Committees. The Atherosclerosis Risk in Communities Study is carried out as a collaborative study supported by National Heart, Lung, and Blood Institute contracts N01-HC-55015, N01-HC-55016, N01-HC-55018, N01-HC-55019, N01-HC-55020, N01-HC-55021, N01-HC-55022, R01HL087641, R01HL59367 and R01HL086694; National Human Genome Research Institute contract U01HG004402; and National Institutes of Health contract HHSN268200625226C. LABJ was supported by a Competitiveness Operational Program Grant of the Romanian Ministry of European Funds (HINT, ID P_37_762; MySMIS 103587).

The authors thank the staff and participants of the ARIC study for their important contributions. Infrastructure was partly supported by Grant Number UL1RR025005, a component of the National Institutes of Health and NIH Roadmap for Medical Research. The Framingham Heart Study and the Framingham SHARe project are conducted and supported by the National Heart, Lung, and Blood Institute (NHLBI) in collaboration with Boston University. The Framingham SHARe data used for the analyses described in this manuscript were obtained through dbGaP. This manuscript was not prepared in collaboration with investigators of the Framingham Heart Study and does not necessarily reflect the opinions or views of the Framingham Heart Study, Boston University, or the NHLBI.

---

**Supplemental Table 1.** Characteristics of sample sets

|  | European (N=16049) |  |  | Combined Polynesian (N=1608) |  |  |
| --- | --- | --- | --- | --- | --- | --- |
|  | Gout | HU | NU | Gout | HU | NU |
| N | 1699 | 2422 | 11928 | 912 | 202 | 494 |
| N Male (%) | 1439 (84.7) | 1909 (78.8) | 4773 (40.0) | 750 (82.2) | 147 (72.8) | 177 (35.8) |
| Age at recruitment, yrs – mean (sd) | 62.99 (12.9) | 54.79 (12.7) | 53.35 (12.9) | 53.22 (13.2) | 43.21 (15.4) | 44.86 (14.6) |
| Urate at recruitment, mmol/l - mean (sd) | 0.41 (0.13) [85.2]* | 0.47 (0.05) | 0.3 (0.07) | 0.43 (0.12) [90.6] | 0.48 (0.05) | 0.33 (0.06) |
| Highest recorded urate, mmol/l - mean (sd)^ | 0.48 (0.14) [95.5] | 0.47 (0.05) | 0.3 (0.07) | 0.54 (0.12) [95.3] | 0.49 (0.06) | 0.34 (0.06) |
| BMI, kg/m<sup>2</sup> - mean (sd) | 30.34 (7.09) [87.5] | 29.13 (4.93) [80.6] | 26.25 (4.72) [81.3] | 36.31 (8.07) [94.4] | 35.46 (7.34) [98.0] | 31.89 (7.09) [96.6] |
| Age of gout onset, yrs - mean (sd) | 47.6 (15.9) [85.8] | - | - | 38.80 (14.38) [94.4] | - | - |
| N attacks in past year - mean (sd) | 4.52 (8.59) [54.9] | - | - | 6.87 (10.75) [92.1] | - | - |
| N *rs2231142* genotypes available (%) | 1637 (96.4) | 2378 (98.2) | 11604 (97.3) | 912 (100) | 202 (100) | 494 (100) |
| N *rs10011796* genotypes available (%) | 1580 (93.0) | 2422 (100) | 11927 (100) | 910 (99.8) | 202 (100) | 494 (100) |
| N *rs11942223* genotypes available (%) | 1576 (92.8) | 2422 (100) | 11927 (100) | 912 (100) | 202 (100) | 494 (100) |

\*All individuals had age and sex data at a minimum. All HU and NU controls had urate at recruitment. Some gout patients did not have urate at recruitment. Figures in italic square brackets represent the proportion of the sample set for which data were available.
^Highest recorded urate is the maximum urate value available in the data across variables pre-ULT urate, urate at diagnosis, urate at recruitment, urate in medical records. Individuals may have none (gout patients only), just one or all of these urate measures.

**Supplemental Table 2.** Association analysis of *rs2231142, rs10011796 (ABCG2)* and *rs11942223 (SLC2A9)* in European and NZ Polynesian sample sets with the risk of gout in males only

| SNP | Raw N | Genotype, N (%)<sup>†</sup> |  |  | Effect allele, N (%) | Gout vs. HU*<sup>^</sup><br>OR (95% CI), *P* | Gout vs. All Controls*<br>OR (95% CI), *P* | HU vs. NU*<br>OR, 95% CI, *P* |
| --- | --- | --- | --- | --- | --- | --- | --- | --- |
| ***rs2231142*** |  | **GG** | **GT** | **TT** | **T** |  |  |  |
| European gout | 1388 | 843 (60.7) | 475 (34.2) | 70 (5.0) | 615 (22.2) | 1.91 (1.67-2.20), *4.7E-20* |  | - |
| European HU | 1872 | 1398 (74.7) | 443 (23.7) | 31 (1.7) | 505 (13.5) | 1.96 (1.71-2.26), *4.8E-21* | 2.53 (2.26-2.84), *1.6E-58* | 1.60 (1.42-1.80), *7.3E-15* |
| European NU | 4642 | 3861 (83.2) | 739 (15.9) | 42 (0.9) | 823 (8.9) | - |  |  |
| WP gout | 330 | 83 (25.2) | 170 (51.5) | 77 (23.3) | 324 (49.1) | 1.95 (1.23- 3.17), *5.4E-03* |  | - |
| WP HU | 66 | 32 (48.5) | 24 (36.4) | 10 (15.2) | 44 (33.3) | 1.81 (1.10-3.03), *0.022* | 2.60 (1.81-3.81), *4.3E-07* | 2.10 (1.15-4.01), *0.020* |
| WP NU | 68 | 40 (58.8) | 24 (35.3) | 4 (5.9) | 32 (23.5) | - |  |  |
| EP gout | 420 | 333 (79.3) | 86 (20.5) | 1 (0.2) | 88 (10.5) | 2.86 (1.28-7.41), *0.017* |  | - |
| EP HU | 81 | 74 (91.4) | 7 (8.6) | 0 (0.0) | 7 (4.3) | 3.03 (1.32-8.00) *0.015* | 3.56 (1.95-6.94), *7.8E-05* | 1.38 (0.41- 4.64), *0.59* |
| EP NU | 109 | 102 (93.6) | 7 (6.4) | 0 (0.0) | 7 (3.2) | - |  |  |
| ***rs10011796*** |  | **CC** | **CT** | **TT** | **T** |  |  |  |
| European gout | 1335 | 278 (20.8) | 646 (48.4) | 411 (30.8) | 1468 (55.0) | 1.43 (1.28-1.58), *3.7E-06* |  | - |
| European HU | 1909 | 525 (27.5) | 945 (49.5) | 439 (23.0) | 1823 (47.7) | 1.42 (1.28-1.59), *3.7E-11* | 1.59 (1.46-1.74), *1.1E-25* |  |
| European NU | 4773 | 1511 (31.7) | 2381 (49.9) | 881 (18.5) | 4143 (43.4) | - | - | 1.20 (1.11-1.29), *3.7E-06* |
| WP gout | 330 | 35 (10.6) | 145 (43.9) | 150 (45.5) | 445 (67.4) | 1.09 (0.70-1.69), *0.69* |  | - |
| WP HU | 66 | 7 (10.6) | 32 (48.5) | 27 (40.9) | 86 (65.2) | 1.03 (0.64-1.66), *0.89* | 1.18 (0.84-1.64), *0.34* |  |
| WP NU | 68 | 8 (11.8) | 34 (50.0) | 26 (38.2) | 86 (63.2) | - |  | 1.12 (0.63-1.99), *0.70* |
| EP gout | 418 | 73 (17.5) | 204 (48.8) | 141 (33.7) | 486 (58.1) | 0.95 (0.65-1.38), *0.78* |  | - |
| EP HU | 81 | 14 (17.3) | 34 (42.0) | 33 (40.7) | 100 (61.7) | 0.89 (0.60-1.31), *0.55* | 0.94 (0.72-1.22), *0.63* |  |
| EP NU | 109 | 19 (17.3) | 49 (45.0) | 41 (37.6) | 131 (60.1) | - |  | 1.06 (0.69-1.63), *0.78* |
| ***rs11942223*** |  | **TT** | **TC** | **CC** | **C** |  |  |  |
| European gout | 1332 | 1168 (74.1) | 382 (24.2) | 26 (1.6) | 373 (14.0) | 0.70 (0.60-0.81), *1.6E-06* | 0.56 (0.50-0.63), *9.9E-21* | - |
| European HU | 1909 | 1658 (68.5) | 701 (28.9) | 63 (2.6) | 694 (18.2) | 0.73 (0.61-0.82), *3.4E-06* |  |  |
| European NU | 4772 | 7101 (59.5) | 4220 (35.4) | 606 (5.1) | 2249 (23.6) | - |  | 0.71 (0.65-0.79), *7.07E-12* |
| WP gout | 330 | 314 (95.2) | 16 (4.9) | 0 (0.0) | 16 (2.4) | 0.92 (0.22-6.31), *0.91* |  | - |
| WP HU | 66 | 64 (97.0) | 2 (3.0) | 0 (0.0) | 2 (1.5) | 0.88 (0.19-6.30), *0.88* | 0.95 (0.33-3.16), *0.93* | 0.54 (0.06-3.71), *0.53* |
| WP NU | 68 | 65 (95.6) | 3 (4.4) | 0 (0.0) | 3 (2.2) | - |  |  |
| EP gout | 420 | 386 (91.9) | 31 (7.4) | 3 (0.7) | 37 (4.4) | 0.97 (0.40-2.78), *0.95* |  | - |
| EP HU | 81 | 74 (91.4) | 7 (8.6) | 0 (0.0) | 7 (4.3) | 1.10 (0.43-3.26), *0.86* | 1.18 (0.61-2.45), *0.63* | 2.56 (0.75-8.92), *0.13* |
| EP NU | 109 | 102 (93.6) | 7 (6.4) | 0 (0.0) | 7 (3.2) | - |  |  |

\*European sample set adjusted by age. Polynesian sample sets additionally adjusted by PCAs 1-10.
^For Gout vs. HU analyses, the top figure is before adjustment by highest recorded serum urate, and the bottom figure is after adjustment.

